# Selective IL13Rα2-Targeted Functionality of IL13-Ligand CARs is Enhanced by Inclusion of 4-1BB Co-Stimulation

**DOI:** 10.1101/2022.03.30.486439

**Authors:** Renate Starr, Xin Yang, Brenda Aguilar, Diana Gumber, Stephanie Huard, Dongrui Wang, Wen-Chung Chang, Alfonso Brito, Vivian Chiu, Julie R. Ostberg, Benham Badie, Stephen J. Forman, Darya Alizadeh, Leo D. Wang, Christine E. Brown

**Affiliations:** Department of Hematology and Hematopoietic Cell Transplantation, T Cell Therapeutics Research Laboratories, Beckman Research Institute, City of Hope National Medical Center, Duarte, CA; Department of Immuno-oncology, Beckman Research Institute, City of Hope National Medical Center, Duarte, CA; Department of Pediatrics, City of Hope National Medical Center, Duarte, CA; Department of Neurosurgery, City of Hope National Medical Center, Duarte, CA; Department of Developmental and Stem Cell Biology, Beckman Research Institute, City of Hope National Medical Center, Duarte, CA

**Keywords:** Chimeric antigen receptor, glioblastoma, costimulatory domains, IL13Rα2, T cell

## Abstract

Chimeric antigen receptor (CAR) T cell immunotherapy is emerging as a powerful strategy for cancer therapy; however, an important safety consideration is the potential for off-tumor recognition of normal tissue. This is particularly important as ligand-based CARs are optimized for clinical translation. Our group has developed and clinically translated an IL13(E12Y) ligand-based CAR targeting the cancer antigen IL13Rα2 for treatment of glioblastoma (GBM). There remains limited understanding of how IL13-ligand CAR design impacts the activity and selectivity for the intended tumor-associated target IL13Rα2 versus the more ubiquitous unintended target IL13Rα1. In this study, we functionally compared IL13(E12Y)-CARs incorporating different intracellular signaling domains, including first-generation CD3ζ- containing CARs (IL13ζ), second-generation 4-1BB- (CD137) or CD28-containing CARs (IL13- BBζ or IL13-28ζ), and third-generation CARs containing both 4-1BB and CD28 (IL13-28BBζ). *In vitro* co-culture assays at high tumor burden establish that 2^nd^ generation IL13-BBζ or IL13- 28ζ outperform first-generation IL13ζ and 3^rd^ generation IL13-28BBζ CAR designs, with IL13- BBζ providing superior CAR proliferation and *in vivo* anti-tumor potency in human xenograft mouse models. IL13-28ζ displayed a lower threshold for antigen recognition, resulting in higher off-target IL13Rα1 reactivity both *in vitro* and *in vivo*. Syngeneic mouse models of GBM also demonstrate safety and anti-tumor potency of murine IL13-BBζ CAR T cells delivered systemically after lymphodepletion. These findings support the use of IL13-BBζ CARs for greater selective recognition of IL13Rα2 over IL13Rα1, higher proliferative potential, and superior anti-tumor responsiveness. This study exemplifies the potential of modulating factors outside the antigen targeting domain of a CAR to improve selective tumor recognition..

## INTRODUCTION

Chimeric antigen receptor (CAR) T cell therapy has achieved striking efficacy in the treatment of relapsed and refractory CD19^+^ B cell malignancies (1, 2), and there is tremendous interest in extending these gains into other tumor types, solid tumors in particular. An evolving area of study is the impact of CAR design, e.g., affinity optimization, spacer selection, or choice of intracellular signaling domain, on differential target recognition. A better understanding of these principles will permit the design of CARs with superior specificity and diminished off-target toxicity, as well as improved expansion, persistence, and anti-tumor potency.

Our group has developed CAR T cells that recognize the high-affinity interleukin-13 receptor α2 (IL13Rα2). IL13Rα2 is expressed on both adult and pediatric brain tumors, including high grade glioma, ependymoma, atypical teratoid/rhabdoid tumor, and brainstem glioma (3–5). Importantly, IL13Rα2 is not expressed on normal brain tissue (3–5). Our CAR designs incorporate IL13, the natural ligand for IL13Rα2, as the antigen-binding domain, with an E12Y engineered mutation that improves selectivity of the CAR for IL13Rα2 over IL13Rα1(6–8), which is expressed more frequently on normal tissues. Additionally, based on preclinical and clinical studies, we have incorporated numerous other construct modifications to optimize clinical and biological activity. These include mutations in the hinge and spacer domains that reduce interactions with Fc gamma receptors (FcγRs) (9), as well changes in the cytoplasmic endodomain from a 1st-generation CD3ζ-CAR (IL13-ζ; containing only ITAM motifs) to a 2nd-generation 4-1BB-containing CAR (IL13-BBζ) (6,10–12). Finally, we have also evaluated route of delivery as an important contextual modification that greatly improves CAR T cell efficacy (6). Taken together, this body of work shows that 2^nd^ generation IL13-BBζ CAR T cells are superior to 1^st^ generation CAR T cells in controlling GBM xenotransplants in mice (6), and that intraventricular delivery of IL13- BBζ CAR T cells provides improved control of multifocal disease as compared to intratumoral or intravenous (IV) delivery (6). These studies provide the foundation for several phase I clinical trials for refractory or recurrent brain tumors (NCT02208362, NCT03389230, NCT04003649, NCT04214392, NCT04510051), in which 2^nd^ generation 4-1BB-containing CAR T cells are delivered intracranially to patients.

Expanding on this work, herein we investigate the molecular underpinnings of important clinical observations of the IL13-BBζ CAR constructs and identified further contextual modifications that improve IL13Rα2-CAR function *in vivo*. Specifically, we describe important functional differences between 1^st^, 2^nd^, and 3^rd^ generation IL13Rα2-targeted CAR constructs, identifying 2^nd^ generation CARs as superior to both 1^st^ and 3^rd^ generation constructs *in vitro* and *in vivo*. We show that IL13-28ζ CAR T cells are more likely than IL13-BBζ CAR T cells to cause off-target toxicity through IL13Rα1 recognition, and that IL13-BBζ CAR T cells are more effective than IL13-28ζ CAR T cells at the low effector-to-target (E:T) ratios reflective of the *in vivo* setting.

We use protein-focused techniques to compare signaling upon antigen stimulation between these 2^nd^ generation CAR constructs and find that IL13-BBζ CAR T cells activate the noncanonical NFκB pathway more effectively than IL13-28ζ CAR T cells, particularly in the setting of high antigen concentration. Finally, we show that immunocompetent syngeneic mice treated with systemic administration of IL13-BBζ CAR T cells after lymphodepleting irradiation exhibit no evidence of off-tumor toxicity and mediate potent antitumor activity in murine orthotopic glioma models. Excitingly, lymphodepleted mice that successfully clear disease after CAR T cell treatment are subsequently able to survive rechallenge with antigen-negative tumor, indicating

that they develop immunologic memory to other tumor-associated antigens. These findings, taken together, indicate that IL13-BBζ CAR constructs are more suited for further clinical development than IL13-28ζ CAR constructs, and justify further clinical trials to evaluate the effects of contextual modifications such as lymphodepletion on IL13-BBζ CAR T cell efficacy.

## MATERIALS AND METHODS

### CAR constructs

The codon-optimized IL13(E13Y) mutein-containing, first generation IL13-zetakine CAR sequence was previously described (11), and is here referred to as IL13-ζ. The 4-1BB-containing 2^nd^-generation CAR construct (IL13-BBζ) was also previously described (6), with the CD28- containing 2^nd^ generation CAR (IL13-28ζ) as well as the 3^rd^ generation CAR (IL13-28BBζ) only differing by the co-stimulatory sequences that were inserted by splice overlap PCR (**Fig. 1A**).

**Figure 1.**
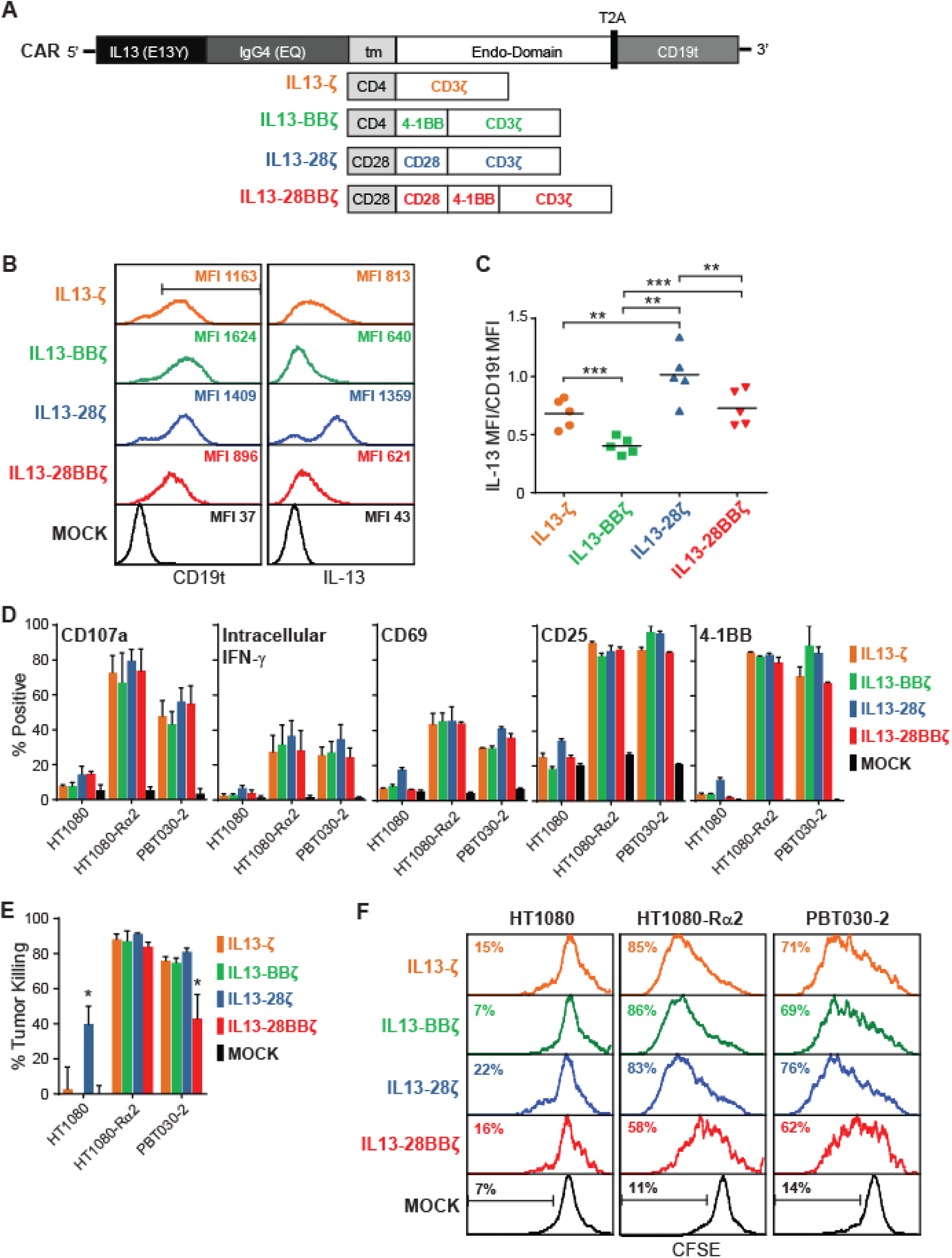
Design, expression, and effector activity of IL13(E12Y)-CAR variants. **A**, Schematics of the cDNA open reading frame of each construct, where the IL13Rα2-targeting ligand IL13(E13Y), IgG4(EQ) Fc hinge, transmembrane (tm), and cytoplasmic endo-domains consisting of the indicated 4-1BB and/or CD28 co-stimulatory and CD3ζ signaling domains of each CAR variant, as well as the T2A ribosomal skip and truncated CD19 (CD19t) sequences are indicated. **B**, Representative flow cytometric assessment of transgene expression on Tcm-derived cells that were mock-transduced (MOCK) or transduced to express the indicated IL13Rα2-CAR variant. Left histograms depict the transduction efficiency based on CD19t expression. The region marker in the top left histogram indicates the CD19t+ gating strategy for the cells displayed in the right histogram, which depicts expression of the IL13-containing IL13Rα2- CAR. **C**, Relative CAR expression level, as defined by the ratio of IL13 to CD19t mean fluorescence intensity (MFI). Data are generated from Tcm-derived cells transduced to express the indicated IL13(E13Y)-CAR variants from five different human donors, expanded and enriched as described in the Methods, and stained as depicted in B. Lines indicate the mean values; using a paired Student’s t-test: **, p < 0.01, ***, p < 0.001. **D,** Characterization of antigen-induced activation. CD107a: Degranulation of IL13Rα2-CAR T cell variants upon challenge with IL13Rα2-negative parental HT1080 fibrosarcoma cells, IL13Rα2-expressing HT1080-Rα2, or PBT030-2 cells for 5 hours at a 1:1 E:T ratio. Mean + S.D. of % CD107a+ in CD3/CD19t+ gated cells from three different donors are depicted. Intracellular IFN-γ: expression in IL13Rα2-CAR T cell variants after 5-hour co-culture with the indicated stimulators at a 1:1 E:T ratio; mean + S.D. of CD3/CD19t+ gated cells from three different donors are depicted. CD69, CD25, 4-1BB: Surface expression of activation markers on IL13Rα2-CAR T cell variants after 48-hour co-culture with the indicated stimulators at a 1:4 E:T ratio; mean + S.E.M. of CD3/CD19t+ gated cells from duplicate wells are depicted; data are representative of IL13Rα2- CAR T cell variants from 4 different donors. **E)** Cytotoxic activity of IL13Rα2-CAR T cell variants after 48-hour co-culture with the indicated targets at a 1:4 E:T ratio. Mean + S.D. of triplicate wells are depicted. Data are representative of IL13Rα2-CAR T cells from 4 different donors. Using an unpaired Student’s t-test: *, p < 0.05 when compared to the killing of that same tumor line by each of the other IL13Rα2-CAR T cell variants. **F**) Proliferation of CFSE-stained IL13Rα2-CAR T cell variants after 4 days of co-culture with the indicated stimulators at a 1:1 E:T ratio. CD45/CD19t+ gated cells were analyzed for CFSE dilution (mock-transduced were only gated on CD45+). Data are representative of IL13Rα2-CAR T cell variants from 2 different donors.

### Generation of CAR T cells

PBMC were collected from discard apheresis kits of five different human donors as approved by City of Hope Internal Review Board oversight. Central memory T cells (Tcm) were then selected from the PBMC, transduced with lentivirus to express either of the four CAR variants (**Fig. 1A**), and expanded with IL-2 and IL-15 as previously described (6). The CD3/28 stimulation beads (Fisher Scientific Cat#11141D) were removed on day 7, successfully transduced cells were enriched based on CD19t expression on day 11-15 using EasySep™ Human CD19 Positive Selection Kit II (Stemcell Technologies, Inc., Cat#17854), and the resulting CAR+ T cells were cryopreserved at day 15 or 17. Unless otherwise indicated, thawed cells were then rested overnight in media with IL-2/IL-15 prior to analysis or use in the assays.

Murine IL13-BBζ T cells were generated as previously described (13). Briefly, T cells isolated from mouse spleens, transduced with retrovirus to express the CAR, and expanded with IL-2 and IL-7. Before their use in *in vivo* experiments, the CD3/28 stimulation beads (Invitrogen Cat#11452D) were magnetically separated from the IL13-BBζ T cells, and CAR expression was determined by flow cytometry.

### Cell lines

The patient-derived, low passage GBM tumor sphere line PBT030-2, and PBT030-2 engineered to express the firefly luciferase (ffLuc) reporter gene have been previously described (14). The fibrosarcoma line HT1080, and HT1080 engineered to express IL13Rα2 have also been previously described (6). Jurkat T cells were obtained from the American Type Culture Collection (ATCC #TIB-152) and maintained between 1 x 10^5^ and 1 x 10^6^ in RPMI 1640 (Corning) with 10% FBS (Genesee Scientific), Penicillin-Streptomycin (Gibco), and L-glutamine (Corning). Small cell lung carcinoma line A549 (ATCC #CCL-185) was cultured under ATCC suggested conditions. Mouse GBM lines KLuc and KLuc-IL13Rα2 have been previously described (13).

### Flow cytometry

IL13Rα2-CAR T cell variants were stained with fluorochrome-conjugated monoclonal antibodies (mAbs) to human IL13 (BD Biosciences Cat#340508) and CD19 (BD Biosciences Cat#557835). A549 cells were stained for human IL13Rα1 and IL13Rα2 using R&D Systems goat polyclonal reagents Cat#AF152 and Cat#AF146, respectively, with phycoerythrin-conjugated donkey anti-goat secondary antibody (Novus Biologicals Cat#NB7590). Isotype-matched mAbs served as controls, and DAPI (Life Technologies) was used to determine viability. Data acquisition was performed on a MACSQuant (Miltenyi Biotec) using either FCS Express (De Novo Software) or FlowJo (v10, TreeStar) software.

### In vitro co-culture assays

For degranulation and intracellular IFN-γ assays, T cells and tumor cells were co-cultured at a 1:1 effector to target (E:T) ratio for 5 hours in the presence of GolgiStop Protein Transport Inhibitor (BD Biosciences). Surface phenotype of cells was determined by flow cytometry using fluorochrome-conjugated antibodies specific for CD3 (Miltenyi Biotec, Inc. Cat#130-094-363), CD19 and CD107a (BD Biosciences Cat#555800), followed by permeabilization with Cytofix/cytoperm kit solution (BD Biosciences) and staining with anti-IFN-γ (BD Biosciences Cat#554702). For cytotoxicity and surface expression of activation markers, T cells and tumor cells were co-cultured at a 1:4 E:T ratio for 48 hours, and analyzed by flow cytometry with DAPI using antibodies specific for CD45, CD3, CD19, CD25, CD69, and 4-1BB (BD Biosciences Cat#s 347464, 563109, 557835, 555431, 340560, 555956). Percentages of tumor killing were based on viable tumor cells (CD45-negative, DAPI-negative) in co-cultures with mock-transduced T cells. Long term killing assays were carried out with co-cultures at a 1:20 E:T ratio (1,250:25,000 in a 96-well plate) for 9 or 14 days, and included staining with antibody specific for CD4 (BD Biosciences Cat#562970) where indicated. Proliferation assays used the CellTrace™ CFSE Cell Proliferation kit (Invitrogen) to stain the T cells prior to their co-culture with tumor cells at a 1:1 E:T ratio for 4 days. Cells were then stained with antibodies specific for CD45 and CD19 prior to flow cytometric analysis. Rechallenge assays were carried out as previously described (15), with cells analyzed by flow cytometry using DAPI and antibodies specific for IL13, CD45, CD4, and CD8 (Fisher Scientific Cat#BDB348793) at each time point. Flow cytometric evaluation of intracellular cleaved caspase-3 was carried out using the FITC Active Caspase-3 Kit (BD Bioscience) and CD19 staining after 24 hour co-culture at a 1:4 E:T ratio.

### Bead-bound receptor stimulation assay

Protein G Dynabeads (Invitrogen) were washed and resuspended in PBS with 0.02% Tween-20 (PBS-T) and loaded with recombinant human IL13Rα2-Fc chimera (R&D Systems Cat#614INS; 2.1 µg per 3.9×10^7^ beads) by incubation for 10 minutes at room temperature. After washing in PBS-T, the IL13Rα2-Fc-bound beads were resuspended at 3.90×10^7^ beads/100uL of warmed media. CAR+ T cells (Jurkat- or human donor-derived) were plated at 2×10^5^ cells/well on a 96- well U-bottom plate with or without beads at a 195:1 bead-to-cell ratio and incubated for 16 (Jurkat T cells) or 12 (human donor derived T cells) hours at 37°C. After incubation, T cells were harvested, washed with ice-cold PBS, and resuspended in ice-cold lysis buffer containing a working dilution of Halt Phosphatase Inhibitor Single-Use Cocktail (ThermoFisher). After 30 minutes of incubation on ice, lysates were centrifuged at 17,200*g* for 20 minutes at 4°C, and supernatants were collected and either frozen at -80°C or immediately analyzed by Western blot. In brief, after determining lysate protein concentration by Bradford protein assay, equal proportions of protein were combined with Laemmli buffer (BioRad) and DTT (Sigma-Aldrich) and boiled at 95°C for 5 minutes. Protein was loaded into a 7.5% TGX gel (BioRad) using a Mini-PROTEAN Tetra Cell (BioRad) and transferred to 0.2μm nitrocellulose (Prometheus). Membranes were incubated in blocking buffer for 1 hour at room temperature, washed in TBS with 0.05% Tween-20 (TBST), and then incubated overnight at 4°C with 1:1000 dilution of primary antibody - either anti-p52/p100 (Millipore Sigma Cat#05361) or anti-β-Actin (Cell Signaling Technology Cat#3700). Membranes were then washed and incubated in blocking buffer containing HRP-linked horse anti-mouse (1:5000; Cell Signaling Technology Cat#7076) for 45 minutes at room temperature. After washing with TBST, membranes were imaged on a ChemiDoc Imaging System (BioRad) with SuperSignal Chemiluminescent Substrate (Thermo Scientific).

### Plate-bound receptor assays

T cells were cultured overnight at 5 x 10^3^ cells/well on 96-well plates that had been coated with 5000, 2500, 1250, 625 or 312.5 ng/mL recombinant human IL13Rα1-Fc chimera (R&D Systems Cat#146IR) or IL13Rα2-Fc chimera. Supernatants were then evaluated for IFN-γ levels using the Legend Max ELISA kit with pre-coated plates (human IFN-γ) (Biolegend). Surface phenotype of cells that had been removed from the well was determined by flow cytometry using fluorochrome conjugated antibodies specific for 4-1BB or CD69.

### Mouse studies

All mouse experiments were approved by the City of Hope Institute Animal Care and Use Committee. For xenograft models, on day 0, either ffLuc+ PBT030-2 cells (1×10^5^) were stereotactically implanted into the right forebrain (intracranial, i.c.) of NSG mice (Jackson Laboratory Strain#005557) or A549 cells (1×10^6^) were injected subcutaneously (s.c.) in a 1:1 solution of PBS:Matrigel into the flank of NSG mice. Mice were then treated intratumorally with CAR T cells as indicated in the figure legend for each experiment.

For syngeneic studies, mouse GBM line KLuc-IL13Rα2 cells (1×10^5^) were implanted i.c. in C57BL/6 mice (Jackson Laboratory Strain#000664). Mice were then treated with 500 rads of whole body irradiation on day 5, and/or murine T cells expressing a murine IL13-BBζ CAR (5×10^6^) administered intravenously (i.v.) on day 7. Confirmation of lymphodepletion was performed by flow cytometric analysis of peripheral (retroorbital) blood collected on days 6 and 11. After 110 days, surviving mice were rechallenged i.c. with parental KLuc cells (1×10^4^), and survival was compared to that of naïve mice implanted i.c. with KLuc cells.

Groups of mice were monitored for i.c. tumor engraftment by non-invasive optical imaging as previously described(8) using a Lago-X (Spectral Instruments Imaging), or for s.c. tumor size using calipers. Where indicated, survival was monitored with euthanasia applied according to the American Veterinary Medical Association Guidelines.

### Statistics

Student’s t-tests, ANOVA tests and log rank (Mantel Cox) tests were used as indicated in each figure legend. *, p < 0.05; **, p < 0.01; ***, p < 0.001; ****, p < 0.0001 unless otherwise indicated in the legend.

### Data Availability

Data were generated by the authors and either included in the article or available on request.

## RESULTS

### *In vitro* effector function of IL13(E12Y)-CAR variants with low tumor burden is independent of co-stimulatory domain

We have previously described the preclinical generation and clinical evaluation of T cells expressing IL13Rα2-targeting, ligand-based CARs harboring an IL13 binding domain that contains the E12Y [IL13(E12Y)] single point mutation to minimize cross-reactivity with IL13Rα1 (6-8,10,11). To further reveal the impact of IL13-ligand CAR design on functionality and selectivity, we generated a panel of IL13(E12Y)-CARs harboring different intracellular signaling domains: a 1^st^ generation CD3ζ construct (IL13-ζ), 2^nd^ generation constructs bearing either 4-1BB or CD28 co-stimulatory domains (IL13-BBζ or IL13-28ζ), and a 3^rd^ generation construct containing both 4-1BB and CD28 intracellular signaling domains (IL13-28BBζ) (**Fig. 1A**). Using our well-described manufacturing platform(16), central memory T cells (Tcm) from five different normal human donors were lentivirally transduced to express either one of these IL13(E12Y)-CAR variants, expanded with IL-2 and IL-15, enriched based on expression of the coordinately expressed CD19t transduction marker, and cryopreserved. Interestingly, although the resulting IL13(E12Y)-CAR T cell variants expressed similar levels of CD19t on their cell surface, they expressed different cell surface CAR levels (**Fig. 1B**). In particular, the ratio of CAR to CD19t marker was markedly different between constructs (**Fig. 1C**): IL13-28ζ expressed the highest level of CAR and CAR:CD19t ratio, IL13-BBζ expressed the least CAR relative to CD19t, and IL13-ζ and IL13-28BBζ had intermediate CAR:CD19t ratios.

To test the effects of the different signaling domains on CAR effector activity, we co-cultured the IL13(E12Y)-CAR T cell variants with IL13Rα2-negative HT1080 human fibrosarcoma cells, IL13Rα2-engineered HT1080 (HT1080-Rα2), or the patient-derived GBM line PBT030-2, with glioma stem cell-like characteristics (6). All of the CAR T cell variants, independent of co-stimulatory domain and CAR expression levels, exhibited comparable IL13Rα2-specific degranulation, intracellular IFN-γ production, and activation marker expression at relatively high E:T ratios (1:1, 1:4) (**Fig. 1D**). Each of the IL13(E12Y)-CAR T cell variants were also able to efficiently kill IL13Rα2-expressing tumor cells in a 48-hour co-culture at E:T ratios of 1:4 (**Fig. 1E**), albeit the 3^rd^ generation IL13-28BBζ CAR displayed slightly reduced killing potency against the GBM line PBT030-2. Additionally, all CAR variants robustly proliferated after four days of co-culture with the HT1080-Rα2 and PBT030-2 stimulator lines compared with Mock T cell controls (**Fig. 1F**). IL13-28BBζ T cells, however, did not proliferate as well as the other CAR T cell variants, which may relate to the slightly sub-optimal killing of PBT030-2 that had been observed with this construct (**Fig. 1E**). We also noted that IL13-28ζ appeared to direct higher levels of killing of the IL13Rα2-negative HT1080 cells, and these CAR T cells exhibited higher levels proliferation upon HT1080 stimulation (**Fig. 1E, F**). Overall, these studies indicate that IL13Rα2-dependent activation and effector function of IL13(E12Y)-CARs was not strongly influenced by either CAR expression levels or the co-stimulatory domain.

### IL13-BBζ CAR T cells outperform other CAR variants *in vitro* at high tumor burden

It was surprising to us that initial *in vitro* studies showed little difference in effector activity of the IL13(E12Y)-CAR T cell variants (**Fig. 1**), even for IL13-ζ, which lacked co-stimulatory signaling. As a result, we next examined co-cultures with PBT030-2 cells at E:T ratios that were decreased to a more challenging ratio of 1:20 to evaluate the recursive killing potential of CAR variants (**Fig. 2A-D**). In this 14-day *in vitro* stress test, 2^nd^ generation IL13-BBζ and IL13-28ζ outperformed both IL13-ζ and IL13-28BBζ, demonstrating the requirement for optimal co-stimulation for recursive killing potency. IL13-BBζ CAR T cells exhibited clear superiority with regards to CAR T cell persistence and proliferation, with significant numbers of CAR-positive T cells detected in co-cultures after tumor cell elimination (**Fig. 2B, C**). To further compare the recursive killing potential of IL13(E12Y)-CAR T cell variants, a tumor rechallenge assay was performed with the addition of GBM cells every other day (**Fig. 2D**) (15, 17). All IL13(E12Y)- CAR T cell variants mediated effective elimination of GBM cells until day three (i.e., after a single GBM rechallenge, total E:T = 1:12). However, upon additional GBM tumor challenge, IL13-ζ and IL13-28BBζ CAR T cells showed decreased cytotoxicity (**Fig. 2D**). Between the 2^nd^ generation CAR T cell variants, IL13-BBζ again outperformed IL13-28ζ CAR T cells, with more efficient tumor elimination following repetitive tumor addition (**Fig 2D**).

**Figure 2.**
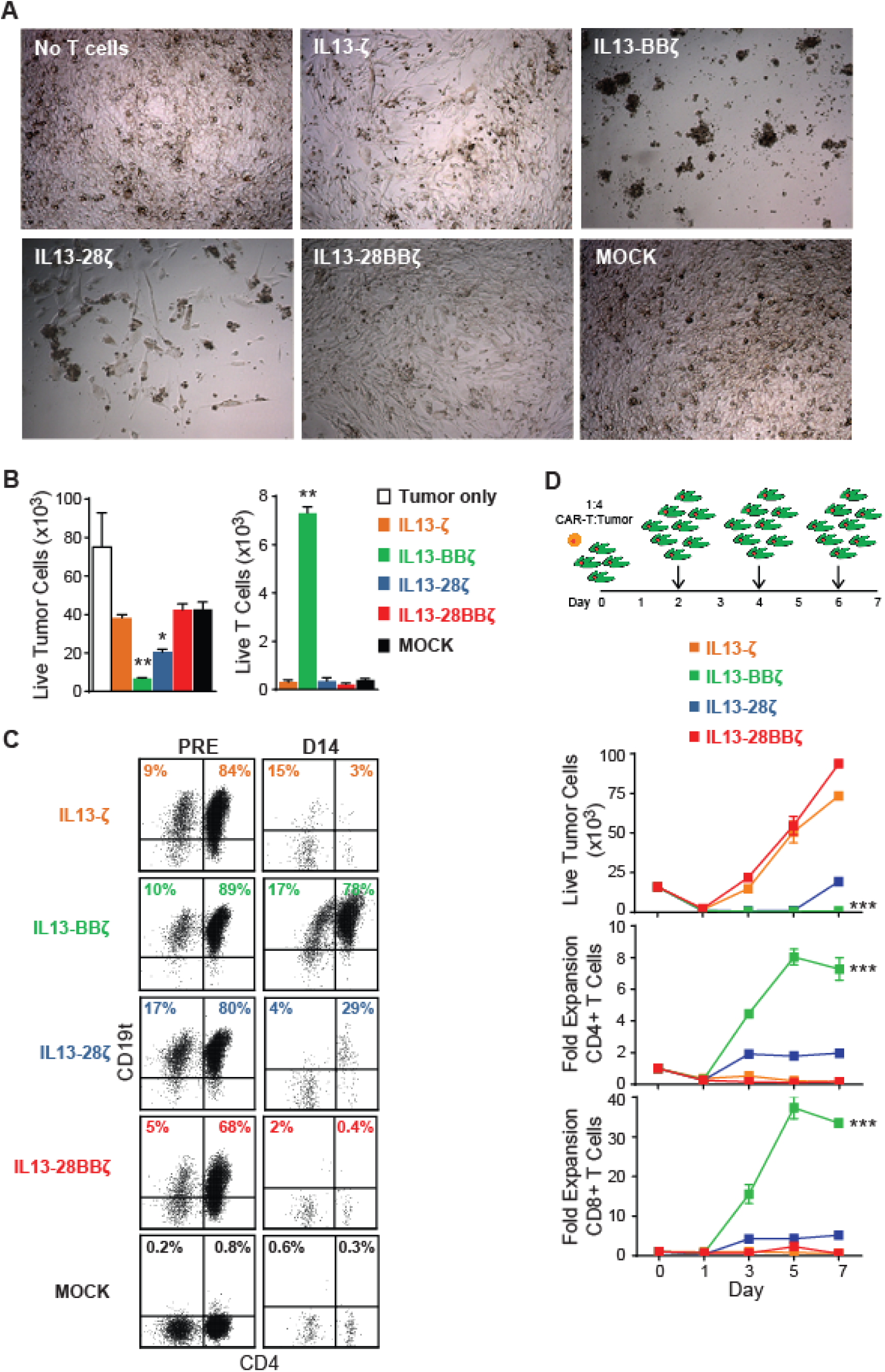
T cells expressing the IL13-BBζ CAR have superior persistence and anti-tumor activity in a long-term co-culture with high tumor burden. **A**) Day 9 culture images of PBT030-2 cells that had been plated alone or co-cultured at a 1:20 E:T ratio with the indicated IL13Rα2-CAR T cell variants. **B**) Enumeration of viable tumor cells (DAPI-/CD45- gated, left) and T cells (DAPI-/CD45+ gated, right) after 14 days of co-cultures that had been plated at a 1:20 E:T ratio as in (A). Mean + S.D. of values from triplicate wells are depicted. Using an unpaired Student’s t-test: *, p = 0.0006; **, p <0.0001 when compared to MOCK. **C**) CD4 and CD19t transgene expression on the IL13Rα2-CAR T cell variants (DAPI-/CD45+ gated) before and after the 14-day co-culture with PBT030-2 as in (B). Percentages of immunoreactive cells are indicated in the upper quadrants, which were drawn based on isotype controls. Data are representative of IL13Rα2-CAR T cell variants from 2 different donors. **D**) The indicated IL13Rα2-CAR T cell variants (4×10^3^ cells) were co-cultured with PBT030-2 cells (1.6×10^4^ cells) and re-challenged with 3.2 ×10^4^ GBM cells every other day (schema at top). Remaining viable tumor cell numbers and fold expansion of either CD4-gated (middle) or CD8-gated (bottom) IL13Rα2-CAR T cells were determined at the indicated time points during the rechallenge assay. Mean ± S.E.M. of values from duplicate wells are depicted. Using a two-way ANOVA test: ***, p < 0.0001 when compared to each of the other IL13Rα2-CAR T cell variants.

To understand whether the differences in persistence may be due to CAR-mediated activation induced cell death (AICD), we evaluated signaling in CAR T cells after antigen stimulation. In CAR T cells co-cultured with PBT030-2 (E:T 1:4) for 24 hours, cleaved caspase-3 was most highly expressed in IL13-28BBζ CAR T cells, whereas IL13-BBζ CAR T cells had the lowest levels, suggesting that IL13-BBζ CAR T cells were the least inclined toward AICD (**Fig. 3A**). It has previously been shown that noncanonical NFκB signaling is upregulated in 4-1BB- containing CAR T cells, and that this leads to increased survival and proliferation (18). First using a Jurkat CAR T model system, we found that IL13-BBζ CAR T cells stimulated for 16 hours with antigen-coated beads upregulated noncanonical NFκB pathway signaling much more than IL13-28ζ CAR T cells as evidenced by an increase in p100 processing (**Fig. 3B, C**).

**Figure 3.**
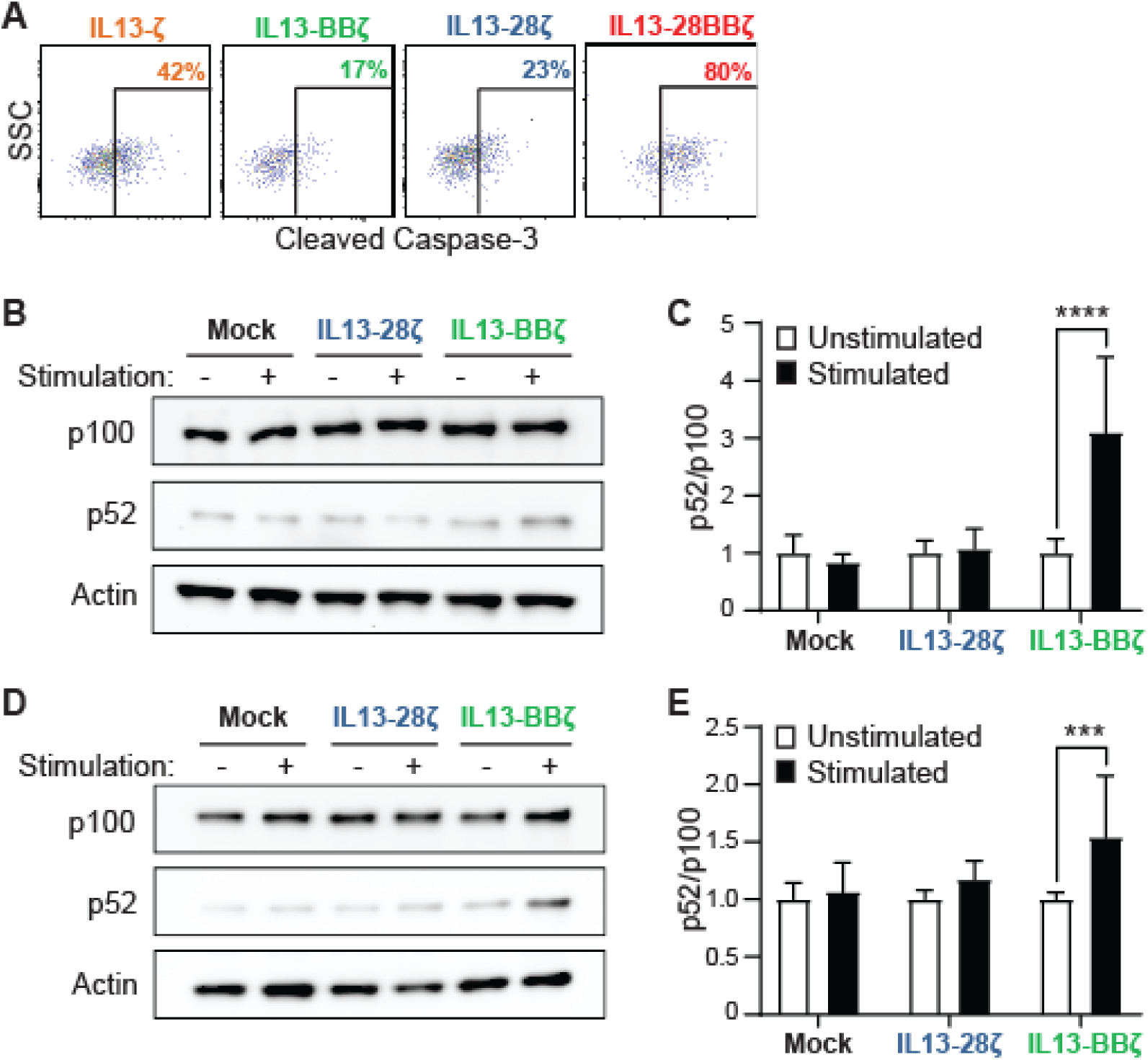
T cells expressing the IL13-BBζ CAR have a greater CAR-induced activation of the ncNF-κB. **A**) The indicated IL13Rα2-CAR T cell variants were co-cultured with PBT030-2 cells at an E:T ratio of 1:4 for 24 hours before they were stained for intracellular cleaved caspase-3. Percentages of immunoreactive CD19t+ gated cells are depicted in each histogram. Using Jurkat (**B, C**) or human donor derived (**D, E**) CAR T cells: **B, D**) Representative Western blot of p52 and p100 abundance in Mock, IL13-28ζ, and IL13-BBζ T cells after overnight IL13Rα2-bead stimulation; and **C, E)** Quantitative analysis of p100 processing, where the band intensity of p52 was divided by the band intensity of p100 for each lane on a Western blot. **C, E**) Mean + S.D. of the ratios of p52 to p100 abundance of 3 different experiments, each using triplicate lanes and normalized to their unstimulated Mock controls, are depicted. Using 2-way ANOVA with Šídák’s multiple comparisons test: ****, p = <0.0001; ***, p = 0.0003 when compared to the relative unstimulated cells. **D, E**) Results representative of CAR T cells derived from 2 different human donors.

Similarly, in CAR T cells generated from a normal human donor, the noncanonical NFκB pathway activation was increased in IL13-BBζ CAR T cells relative to IL13-28ζ CAR T cells after 12 hours after stimulation (**Fig. 3D, E**). Taken together, these findings suggest that the pronounced functional superiority of IL13-BBζ CAR T cells observed at low E:T ratios are a result of improved CAR T cell proliferation and survival through augmented noncanonical NFκB signaling and consequent decreased caspase-3 activity.

### IL13-28ζ CAR T cell activation is more sensitive to lower levels of IL13Rα2 antigen

To evaluate further the impact of antigen density on the functional activity of IL13(E12Y)-CAR T cells, we challenged CAR variants with plate-bound IL13Rα2 at different concentrations (**Fig. 4**). The IL13-BBζ CAR T cells, which functioned well at the low E:T ratios in long-term co- cultures (**Fig. 2**), released IFN-γ levels in a manner similar to the 1^st^ and 3^rd^ generation variants (**Fig. 4A**). This is in contrast to the IL13-28ζ CAR T cells, which produced detectable IFN-γ at antigen concentrations that were 4-fold lower than that observed with any of the other variants (**Fig. 4A**). Analysis of the activation markers 4-1BB and CD69 also suggested that the IL13-28ζ CAR T cells had a lower IL13Rα2-induced activation threshold compared to the IL13-BBζ CAR T cells (**Fig. 4B**).

**Figure 4.**
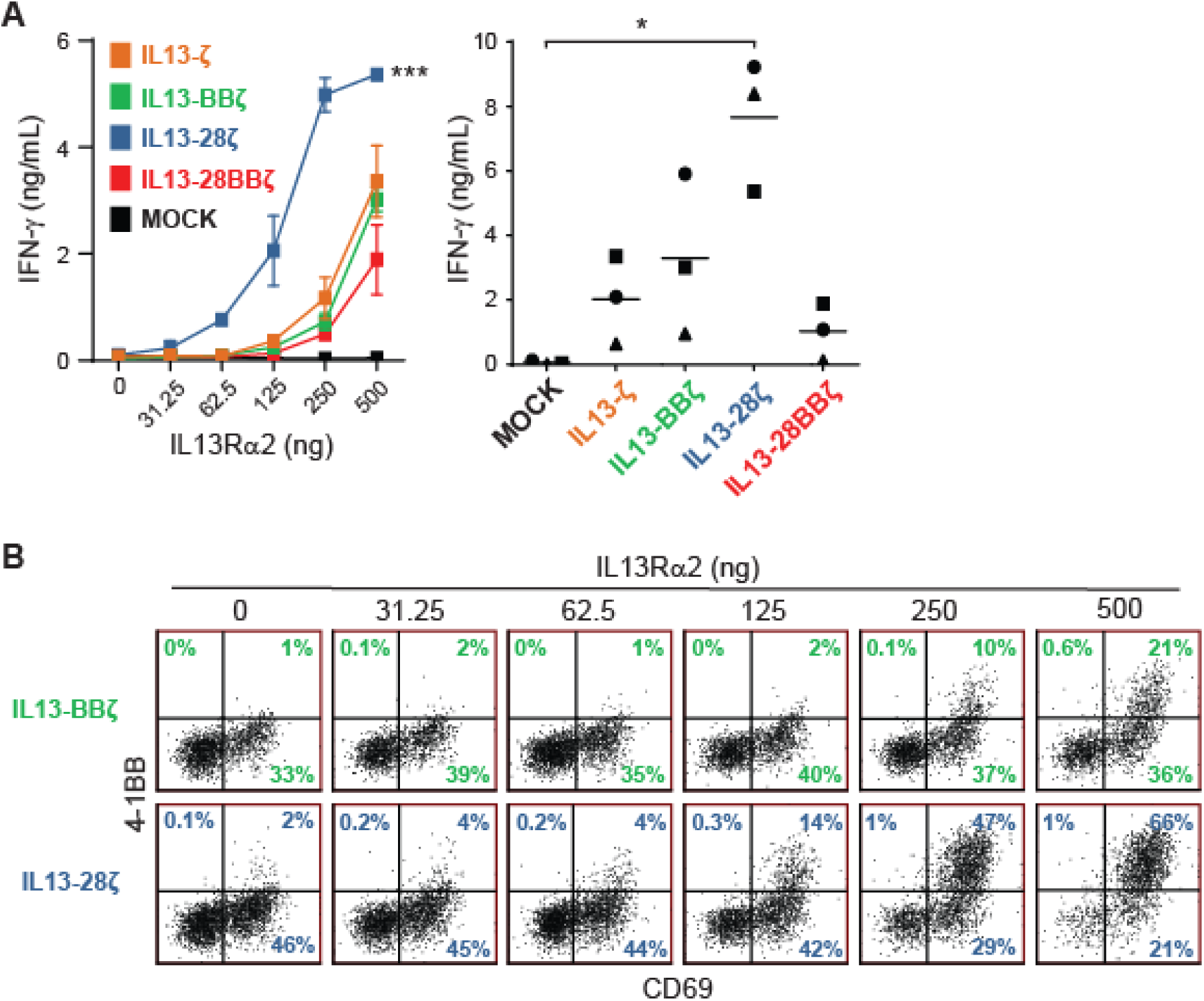
T cells expressing the IL13-28ζ CAR are more sensitive to lower levels of IL13Rα2. **A**, IFN-γ release upon overnight culture of the indicated IL13Rα2-CAR T cell variants on plate-bound IL13Rα2 as determined by ELISA. Left, a representative assay with CAR T cells from a single donor on 0-500 ng plate-bound IL13Rα2, depicting mean ± S.D. of triplicate wells. Using a two-way ANOVA test: ***, p < 0.0001 when compared to each of the other IL13Rα2-CAR T cell variants. Right, results of the indicated IL13Rα2-CAR T cell variants from three different donors cultured on 500 ng plate-bound IL13Rα2, with lines indicating mean values. Using a paired Student’s t-test: *, p = 0.0225. **B**, Surface expression of activation markers on IL13-BBζ and IL13-28ζ CAR T cells from a plate-bound IL13Rα2 assay as in (A) was determined by flow cytometry of the viable CD3/CD19t+ gated cells. Percentages of immunoreactive cells are depicted in each histogram.

### IL13-BBζ CAR T cells exhibit superior IL13Rα2-directed anti-tumor efficacy *in vivo*

We hypothesized that superior persistence and anti-tumor efficacy of IL13-BBζ CAR T cells observed at the low E:T ratios in long-term co-cultures would translate to improved disease control *in vivo*. To test this, we established brain tumor xenografts using ffLuc-expressing PBT030-2, treated the mice intracranially (i.c.) with the IL13Rα2-CAR T cell variants, and then followed tumor size over time using biophotonic imaging (**Fig. 5**). While anti-tumor efficacy could be seen in mice treated with each of the variants, the most significant effects on tumor burden (**Fig. 5B**) and overall survival (**Fig. 5C**) were observed with the IL13-BBζ CAR T cells. Indeed, mice treated with IL13-BBζ CAR T cells exhibited long-term tumor-free survival in >60% of mice after 152 days (**Fig. 5D**).

**Figure 5.**
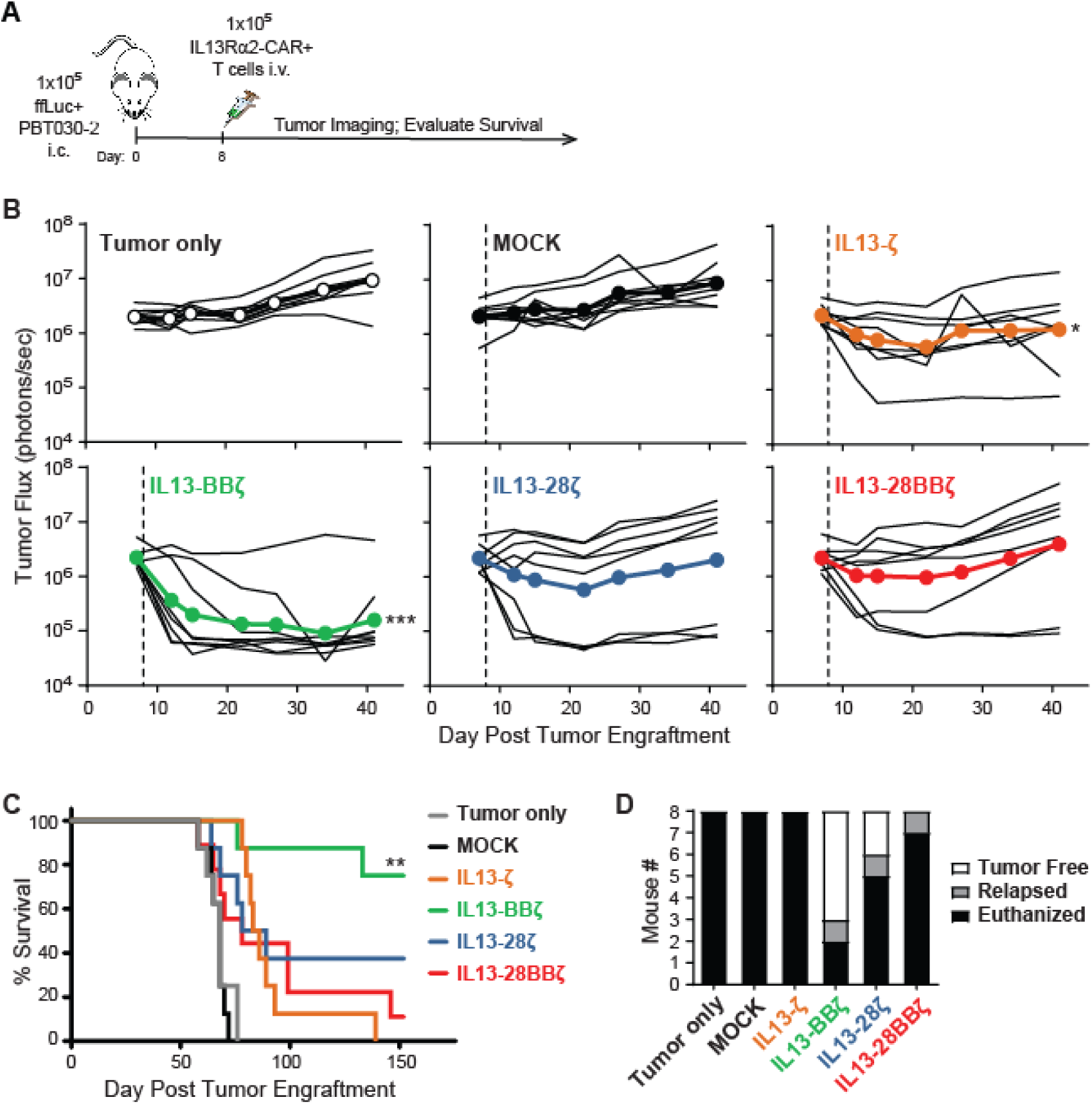
T cells expressing the IL13-BBζ CAR exhibit superior anti-tumor efficacy *in vivo*. A) FfLuc+ PBT030-2 cells (1×10^5^) were implanted i.c. in NSG mice, and 8 days later the indicated IL13Rα2-CAR T cell variants (1×10^5^) were administered intratumorally (n = 8 per group). **B**) Quantification of tumor ffLuc flux (photons/sec) for individual mice over 41 days post tumor engraftment. Dashed lines indicate day of T cell administration. Using a two-way ANOVA test: *, p = 0.0153; ***, p = 0.0002 when compared to MOCK. **C**) Kaplan Meier survival curve (n=8 per group). Using the log rank (Mantel Cox) test: **, p = 0.0021 when compared to IL13-ζ. **D**) Graphic representation of the numbers of euthanized, relapsed, and tumor-free animals in each group at the end point of 152 days.

### IL13-28ζ CAR T cells exhibit off-target activity against IL13Rα1

To further evaluate differences in these IL13Rα2-CAR T cell variants, we examined off-target effects towards IL13Rα1. IL13 binds both IL13Rα1 and IL13Rα2, and while the IL13(E12Y) mutein was originally designed to increase selectivity for IL13Rα2 (7), some groups have reported IL13Rα1 cross reactivity in similar IL13(E12Y)-based CARs containing a CD28 co- stimulatory domain (19, 20). Consistent with these reports, we observed in plate-bound IL13Rα1 assays that our IL13-28ζ CAR produced IFN-γ (**Fig. 6A**) and up-regulated activation marker expression at the highest concentrations of plate bound IL13Rα1-Fc (**Fig. 6B**; 500 ng/mL). By comparison, the other CAR variants, including IL13-BBζ, were not strongly activated at high concentrations of plate-bound IL13Rα1-Fc. Using the tumor line A549, which endogenously expresses IL13Rα1, but not IL13Rα2, we found that the IL13-28ζ CAR T cells exhibited the most off-target-mediated degranulation (**Fig. 6C**), activation marker expression (**Fig. 6D**), *in vitro* cytotoxicity (**Fig. 6E**), and proliferation (**Fig. 6F**). Further, IL13-28ζ CAR T cells exhibited *in vivo* anti-tumor activity against A549 xenograft tumors (**Fig. 6G**). Together, these data argue against using the IL13-28ζ CAR for selective IL13Rα2-targeting of tumors (20).

**Figure 6.**
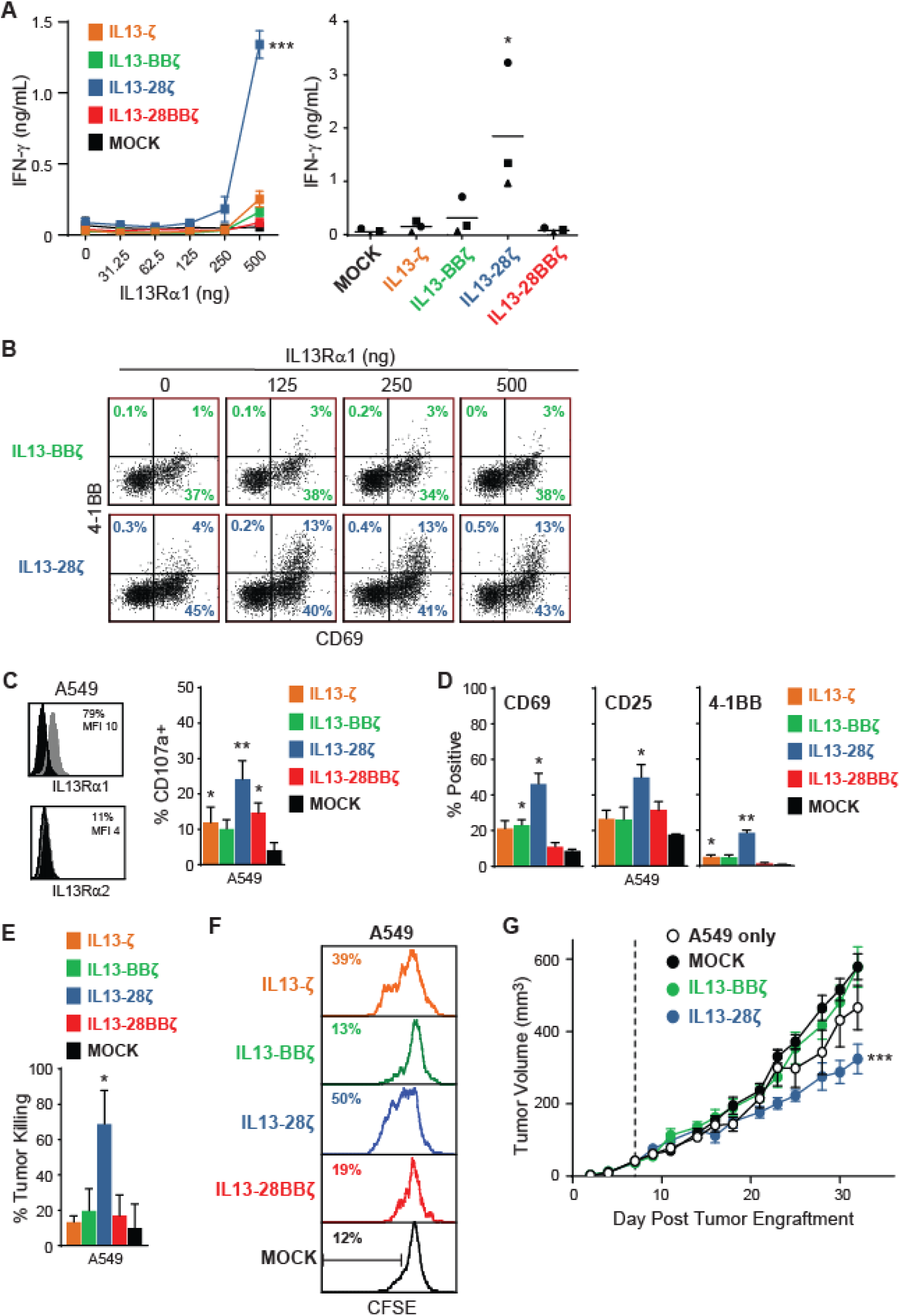
T cells expressing the IL13-28ζ CAR are more cross-reactive to IL13Rα1. **A**) IFN- γ release upon overnight culture of the indicated IL13Rα2-CAR T cell variants on plate-bound IL13Rα1 as determined by ELISA. Left, a representative assay with CAR T cells from a single donor on 0-500ng plate-bound IL13Rα1, depicting mean ± S.D. of triplicate wells. Using a two- way ANOVA test: ***, p < 0.0001 when compared to each of the other IL13Rα2-CAR T cell variants. Right, results of the indicated IL13Rα2-CAR T cell variants from three different donors cultured on 500ng plate-bound IL13Rα1, with lines indicating mean values. Using a paired Student’s t-test: *, p < 0.05 when compared to MOCK. **B**) Surface expression of activation marker CD69 on IL13-BBζ and IL13-28ζ CAR T cells from a plate-bound IL13Rα2 assay as in (A) was determined on viable CD3/CD19t+ gated cells. Percentages of immunoreactive cells are depicted in each histogram. **C**) Degranulation of IL13Rα2-CAR T cell variants upon challenge with IL13Rα1-expressing A549 cells. Mean + S.D. of %CD107a+ of CD3/CD19t+ gated cells from four different donors are depicted. Using a paired Student’s t-test: *, p < 0.05; **, p = 0.006 when compared to MOCK. **D**) Surface expression of activation markers on IL13Rα2-CAR T cell variants after 48-hour co-culture with the indicated stimulators at a 1:4 E:T ratio. Mean + S.E.M. of CD3/CD19t+ gated cells from duplicate wells are depicted. Using an unpaired Student’s t-test: *, p < 0.05; **, p < 0.01 when compared to MOCK. Data are representative of IL13Rα2-CAR T cell variants from 4 different donors. **E**) Cytotoxic activity of IL13Rα2-CAR T cell variants after 48-hour co-culture with A549 targets at a 1:4 E:T ratio. Percentages of tumor killing were based on viable tumor cells in tumor-only wells. Mean + S.D. of triplicate wells are depicted. Using an unpaired Student’s t-test: *, p = 0.012 when compared to MOCK. Data are representative of IL13Rα2-CAR T cells from 4 different donors. **F**) Proliferation of CFSE-stained IL13Rα2-CAR T cell variants after 4 days of co-culture with A549 cells at a 1:1 E:T ratio. CD45/CD19t+ gated cells were analyzed for CFSE dilution (mock-transduced were only gated on CD45+). Data are representative of IL13Rα2-CAR T cell variants from 2 different donors. **G**) A549 cells (1×10^6^) were injected s.c. into the flank of NSG mice and 7 days later the indicated freshly thawed IL13Rα2-CAR T cell variants (2×10^6^) were administered intratumorally. Tumor size was determined with calipers. Dashed line indicates day of T cell administration. Using a two-way ANOVA test: ***, p = 0.0009 when compared to A549 tumor only.

**Figure 7.**
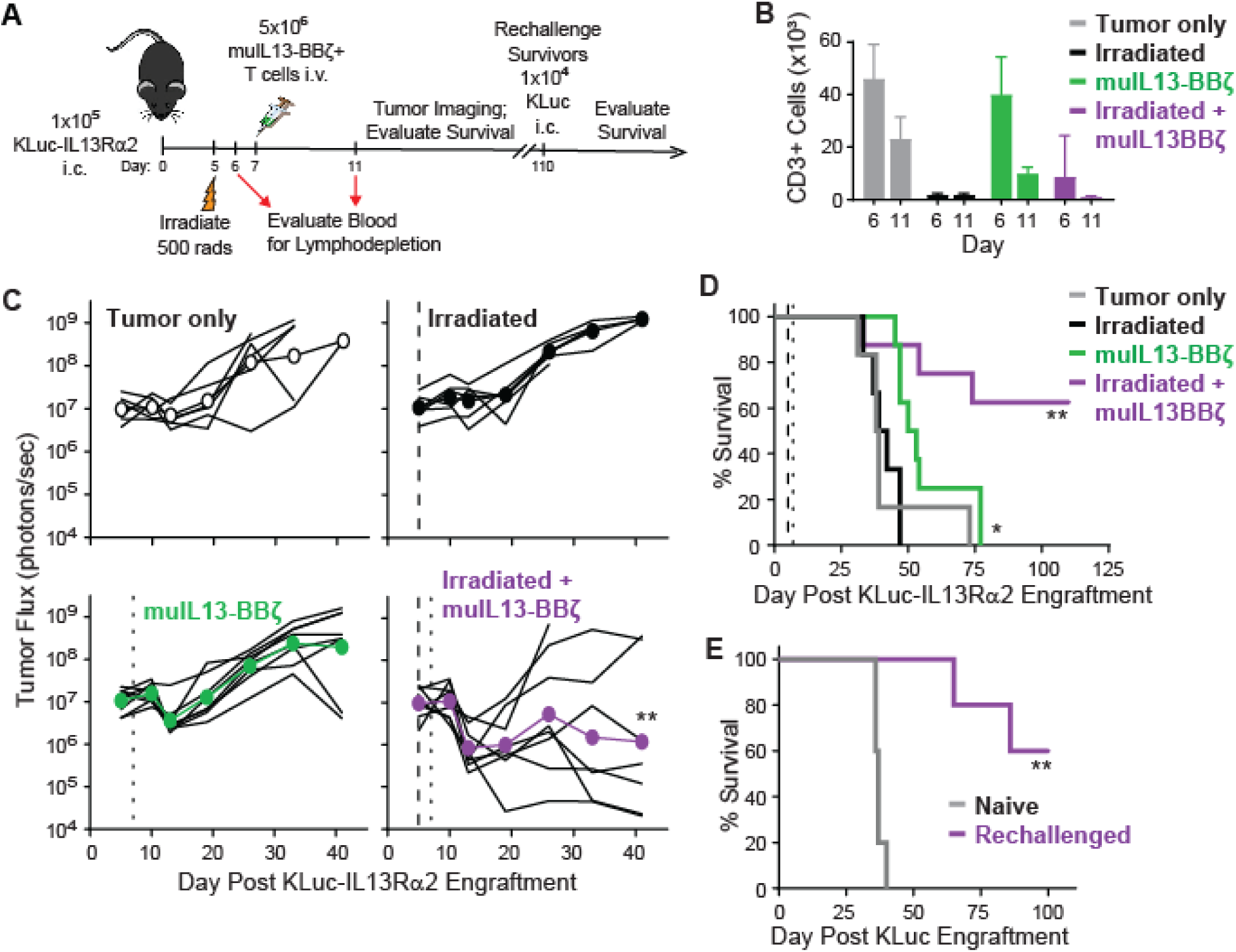
Anti-tumor efficacy of muIL13-BBζ CAR T cells in a syngeneic mouse tumor model. **A**) Mouse GBM line KLuc-IL13Rα2 cells (1×10^5^) were implanted i.c. in C57BL/6 mice, and the indicated groups of mice (n = 6-8 per group) received whole body irradiation on day 5, and/or murine T cells expressing a murine IL13-BBζ CAR (5×10^6^) were administered IV. After 110 days, surviving mice (which occurred only in the irradiated + muIL13-BBζ group) were rechallenged i.c. with parental KLuc cells (1×10^4^), and survival was compared to that of naïve mice implanted i.c. with KLuc cells. **B**) Confirmation of radiation induced lymphodepletion. Enumeration of CD3+ T cells per µL blood was performed on 4 mice per group on days 6 and 11. Means ± S.D. are depicted. **C**) Quantification of tumor luciferase flux (photons/sec) for individual mice over 41 days post tumor engraftment. The dashed line indicates day of irradiation, the dotted line indicates day of T cell administration. Using a mixed model ANOVA test: **, p = 0.0024 when compared to untreated mice. **D**) Kaplan Meier survival curve after engraftment of KLuc-IL13Rα2 cells. Using the log rank (Mantel Cox) test: *, p = 0.0387 and **, p = 0.0048 when compared to untreated mice. **E**) Kaplan Meier survival curve after re-challenge (day 110 depicted here as day 0) with KLuc cells. Using the log rank (Mantel Cox) test: **, p = 0.0019 when compared to naive mice engrafted with KLuc cells for the first time.

### Systemic delivery of murine IL13-BBζ CAR T cells for treatment of GBM is both safe and effective with pre-conditioning lymphodepletion

The above studies comparing human IL13(E12Y)-CAR design support the selective IL13Rα2- targeting of IL13-BBζ CAR T cells. Because *in vitro* co-culture assays and xenograft mouse models likely do not reveal the full potential for off-tumor toxicities given the cross-species differences in IL13-ligand binding, we developed a fully murine IL13(E12Y)-CAR to further study the safety and specific tumor targeting of IL13-BBζ CAR T cells in a syngeneic, immunocompetent mouse brain tumor model (**Fig. 6A**) (13). C57BL/6 splenocytes were engineered to produce muIL13-BBζ CAR T cells that exhibited equivalent numbers of CD4+ and CD8+ subsets with mixed undifferentiated and differentiated T cell populations on day five (13). Brain tumors were established in C57BL/6 mice using KLuc invasive glioma cells derived from *Nf1*, *Trp53* mutant mice(21) that had been further gene modified to express mouse IL13Rα2 (KLuc-IL13Rα2 cells) (13). Since there is interest in clinically evaluating systemic delivery of IL13Rα2-CAR T cells for GBM and other IL13Rα2+ solid tumors, including neuroendocrine cancers (22), melanoma (23), ovarian cancer (24), and colorectal cancer (25), we set out to utilize this fully murine platform to preclinically assess the safety and efficacy of intravenously (i.v.) delivered muIL13-BBζ CAR T cells. Furthermore, because systemic i.v. administration of CAR T cells often incorporates pre-conditioning with lymphodepletion and exacerbates CAR T cell mediated toxicities, we also wanted to evaluate the effect of lymphodepletion on both the safety and anti-tumor potency of the muIL13-BBζ CAR T cells in this model. Using irradiation, we were able to confirm lymphodepletion by flow cytometric analysis of the blood (**Fig. 6B**).

Biophotonic imaging then revealed that the best anti-tumor efficacy was observed in mice treated with both irradiation and muIL13-BBζ CAR T cells (**Fig. 6C**). Overall survival was improved with muIL13-BBζ CAR T cells alone, but the tumors ultimately progressed, and complete cures were only observed upon treatment with both irradiation and muIL13-BBζ CAR T cells (**Fig. 6D**). Furthermore, when the cured mice were re-challenged with i.c. administration of parental KLuc cells (i.e., not expressing muIL13Rα2), 60% survival was again observed, suggesting the establishment in the mice of broader immunological memory against the parental tumor (**Fig. 6E**). Importantly, throughout this study, the mice were monitored for any obvious signs of distress or general toxicity, and those treated with the muIL13-BBζ CAR T cells did not exhibit any signs of therapy-associated adverse events – they did not exhibit weight loss and were bright, alert, and reactive until affected by their tumor burden.

## DISCUSSION

Optimization of CAR T cell therapy for solid tumors faces many challenges; in particular, poor CAR T cell persistence, tumor antigen heterogeneity, antigen escape, and off-tumor targeting present barriers to both the safety and curative potential of CAR T cell therapy. It is increasingly clear that successful therapies will need to combine both rational receptor design and contextual modifications to the tumor microenvironment and immune system. This work with IL13Rα2- CARs advances our understanding of how receptor design impacts CAR expression, signaling, and function. These findings inform our current clinical trials and suggest generalizable strategies for other adoptive cellular therapy efforts.

Our data show that for the IL13Rα2-targeting CAR construct, co-stimulatory domain choice influences cell surface CAR expression. Despite different expression levels, the variants surprisingly conferred comparable recognition, activation and short-term effector activity against target cells *in vitro* (i.e., IL13Rα2-directed degranulation, intracellular IFN-γ production, and activation marker expression). However, in long-term co-cultures with high tumor burden, disparities in persistence, survival and anti-tumor activity of these variants became apparent.

Specifically, IL13-BBζ CAR T cells exhibited superior persistence and recursive killing ability, likely due to increased noncanonical NFκB pathway signaling and decreased caspase-3 signaling. These findings translated into superior IL13Rα2-directed anti-tumor efficacy *in vivo* for the IL13-BBζ variant. Furthermore, IL13-28ζ CAR T cells appeared more sensitive to lower levels of plate-bound IL13Rα2 and IL13Rα1, as evidenced by quantification of 4-1BB and CD69 tumor flux, which showed that IL13-28ζ had a lower activation threshold, and, unlike IL13-BBζ CAR T cells, exhibited off-target activity against IL13Rα1. IL13-28ζ, but not IL13-BBζ, CAR T cells exhibited IL13Rα-dependent degranulation, activation marker expression, cytotoxicity, proliferation, and *in vivo* anti-tumor activity, which is consistent with previous reports evaluating CD28-containing IL13-ligand CAR designs (20). Taken together, these findings argue against using IL13-28ζ for selective IL13Rα2-targeting of tumors in favor of IL13-BBζ CAR T cells.

Other studies have shown CD28-containing CARs to exhibit more robust signaling and lower thresholds of activation than CARs with 4-1BB domains. Indeed, the effect of the costimulatory domain on differential recognition of low antigen density levels has been reported with CARs targeting CD19 (26, 27), HER2(28), PSCA (29), and ROR1 (26). This study extends these findings by showing that costimulatory domains can also impact recognition based on affinity differences between the CAR and its target antigens (i.e., the affinity of IL13(E12Y)-CAR for IL13Rα1 vs. IL13Rα2). Specifically, the CD28 domain increased the off-target recognition of the low affinity IL13Rα1 receptor by the IL13(E12Y)-CAR, whereas that containing the 4-1BB domain demonstrated negligible recognition IL13Rα1 and superior selectivity for IL13Rα2.

To further evaluate the safety of IL13-BBζ CAR T cells, we developed a fully murine IL13(E12Y)-CAR to better assess off-tumor toxicities in immunocompetent mouse models of GBM, and administered the CAR T cells systemically with and without lymphodepletion. Lymphodepletion is known to be required for optimal efficacy with hematologic malignancy- targeted CAR T cells (30, 31) and adoptive therapy of other solid tumors (32); and augments the potency and toxicity profiles of CAR T cell therapy. We have previously shown, both preclinically and clinically, that locoregional delivery of IL13Rα2-CAR T cells was safe and effective at eliminating malignant brain tumors (6, 10). We extend that work here, and, similar to a previous study by Suryadevara et al using lymphodepletion (33), demonstrate that the use of lymphodepleting radiation before i.v. delivery of muIL13-BBζ CAR T cells also significantly enhances *in vivo* anti-tumor efficacy and survival, with complete cures only observed upon treatment with both irradiation and muIL13-BBζ CAR T cells. Importantly, systemic treatment with muIL13-BBζ CAR T cells did not result in any signs of therapy-associated adverse events. Furthermore, when the cured mice were re-challenged, prolonged survival was again observed, suggesting that the establishment of immunological memory in these mice was not compromised by lymphodepleting radiation. Our work contributes to a growing body of evidence supporting the use of pre-conditioning lymphodepletion as a valuable adjunct for solid tumor-directed CAR T cell therapies. Additionally, these studies open the door to investigations combining lymphodepletion and locoregional CAR T cell delivery, studies which are currently ongoing in our group.

Ultimately, this study supports several general principles of CAR design, including improved specificity, persistence, and efficacy of 4-1BB-based 2^nd^ generation CARs, in part, through increased noncanonical NFκB signaling and decreased caspase-3 activity. These differences are most pronounced at lower effector to target ratios, which may help to explain why 4-1BB-based CARs are often clinically superior to CD28-based CARs for solid tumors. Additionally, these findings extend our previous work demonstrating that changes to the context of CAR T cell therapy substantially affect their efficacy; locoregional delivery and lymphodepletion independently augment the ability of IL13-BBζ CAR T cells to control tumors. Taken together, these data justify novel clinical trials that combine IL13-BBζ CAR T cells, locoregional delivery, and lymphodepletion for solid tumor therapy.

## ACKNOWLEDGMENTS

The authors would like to thank Jim O’Hearn and Jennifer K. Shepphird for formatting and editing of this manuscript; and Aniee Sarkissian, Sarah Wright and Brenda Chang for their technical assistance.

## AUTHOR CONTRIBUTIONS

Conception and design: BB, SJF, LDW and CEB Development of methodology: XY, DW, and W-CC Acquisition of data: RS, XY, BA, DG, SH, DW, and VC Analysis and interpretation of data: RS, XY, BA, DG, DW and CEB Writing, review and/or revision of manuscript: DG, JRO, LDW and CEB Administrative, technical or material support: RS, XY, BA, W-CC, and AB Study Supervision: DA, LDW and CEB

